# Preimaginal development of *Aedes aegypti* in brackish water produces adult mosquitoes with thicker cuticles and greater insecticide resistance

**DOI:** 10.1101/2024.07.10.602862

**Authors:** Kokila Sivabalakrishnan, Andrew Hemphill, S.H.P. Parakrama Karunaratne, Arunasalam Naguleswaran, Isabel Roditi, Sinnathamby N. Surendran, Ranjan Ramasamy

## Abstract

*Aedes aegypti* and *Aedes albopictus* mosquitoes, the principal vectors of many human arboviral diseases, lay eggs and undergo preimaginal development in fresh water habitats. They have also recently been found to develop in brackish water in coastal areas. Adult females emerging from brackish water-developing *Ae. aegypti* larvae are shown to possess thicker cuticles and greater resistance to common insecticides used against adults (adulticides) compared with fresh water *Ae. aegypti*. These findings are compatible with previous findings showing that brackish water *Ae. aegypti* larvae possess thicker cuticles and greater larvicide resistance. Greater resistance of salinity-tolerant *Ae. aegypti* to adulticides and larvicides is a hitherto unappreciated problem for controlling arboviral diseases and has implications also for other mosquito-borne diseases.

## Introduction

*Aedes aegypti,* the principal global mosquito vector of arboviral diseases such as dengue, chikungunya, yellow fever and Zika, is widely held to oviposit and undergo preimaginal development only in fresh water (FW) [1-3]. Applying larvicides and minimizing preimaginal FW habitats of *Ae. aegypti* and the secondary arboviral vector *Aedes albopictus*, together with space spraying with insecticides against adults (adulticides), are essential measures for controlling arboviral diseases [1-4]. Dengue is the most prevalent arboviral disease in the world. The WHO reported half the global population to be at risk of infection and 6.5 million cases of dengue resulting in 7300 deaths worldwide in 2023 [1]. Dengue-endemic Sri Lanka and its northern peninsular Jaffna district (Fig. 1), recorded 105,049 and 8,261 dengue cases respectively in 2019, with sharp decreases in 2020 and 2021 during the COVID-19-related people movement restrictions, followed by a gradual return to pre-pandemic levels in 2022 [5].

**Fig. 1.**
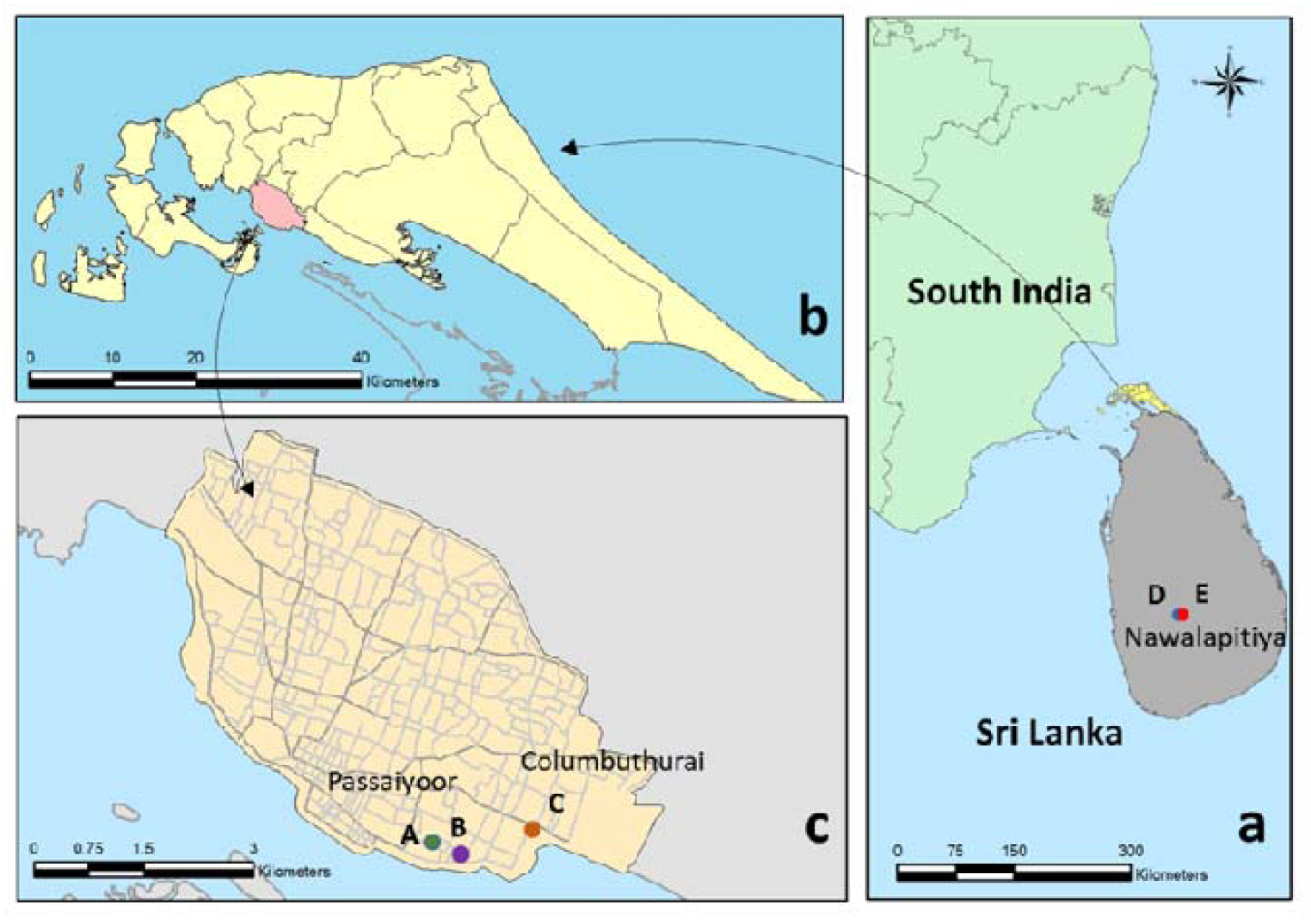
Map of study location and *Aedes aegypti* collection sites. **Legend to Fig. 1 (a)** Location of the island of Sri Lanka and its Jaffna peninsula (highlighted in yellow) in relation to South India. Also shown is Nawalapitiya town in the central hills of Sri Lanka where *Ae. aegypti* larvae were collected from fresh water field habitats at D (7°03’17.8”N 80°32’07.3”E) and E (7°03’19.9”N 80°32’08.8”E); (b) Jaffna peninsula with Jaffna city highlighted in beige; **(c)** Jaffna city showing the sites Passaiyor A (9°38’53.6”N 80°01’49.8”E) and B (9°38’05.4”N 80°01’56.1”E), and Columbuthurai C (9°39’08.5”N 80°02’34.4”E) where *Ae. aegypti* larvae were collected from brackish water field habitats

*Aedes aegypti* and *Ae. albopictus* in the Jaffna peninsula oviposit and undergo preimaginal development not only in FW, but also coastal brackish water (BW) collections of up to 15 g/L salt, *e.g.* in fishing boats, beach debris and coastal wells [6-11]. The ability of both *Aedes* vectors to develop in coastal BW has been confirmed in other countries [11], including the USA [12,13]. FW, BW and saline water are respectively defined as containing <0.5, 0.5 – 30 and >30 g/L salt [6,14]. FW-derived *Ae. aegypti* and *Ae. albopictus* larvae in the Jaffna peninsula are more salinity-tolerant than the corresponding FW-derived larvae from mainland Sri Lanka [6], because of the absence of a barrier to gene flow in *Ae. aegypti* within the small (1100 km^2^) Jaffna peninsula and the peninsula’s relative geographical isolation [11]. Altered cuticle protein composition, increased transcription of genes concerned with cuticle synthesis and cuticle thickening were prominent changes in fourth instar (L4) larvae of BW-adapted *Ae. aegypti* [11,15]. Importantly, the changes in BW *Ae. aegypti* L4 were accompanied by reduced susceptibility to temephos [11], which is the principal larvicide used worldwide for controlling *Aedes* vectors [2-4].

Adaptation of *Ae. aegypti* and *Ae. albopictus,* and others regarded as typical FW mosquitoes, such as the malaria vectors *Anopheles culicifacies* and *Anopheles stephensi*, to develop in BW [8,16,17], heightens the risk of mosquito-borne disease transmission in coastal areas because BW habitats are globally neglected in mosquito vector control programs [2-4,18,19]. Greater disease transmission in coastal areas will spill-over to inland areas [18-20]. Increasing salinization of ground water in coastal zones due to rising sea levels and unsustainable extraction of ground water expand coastal BW habitats for mosquito vectors [18,19], processes that are already evident in the Jaffna peninsula [21]. Increasing salinization in coastal zones is of worldwide concern because about 10% of the global population live in coastal areas [18,19].

A few mosquito vector species found in coastal locations [18,19] oviposit and undergo preimaginal development in brackish water but their cuticle structure and the comparative effectiveness of adulticides for their control are not known. In view of this, and the other context outlined above, it was important to determine whether cuticle structure and susceptibility to common adulticides are altered in adults emerging from preimaginal stages of *Ae. aegypti* developing in BW.

## Results

Experiments were performed with adult female *Ae. aegypti* from (i) self-mating JBW (Jaffna brackish water) and JFW (Jaffna fresh water) laboratory colonies, initially established from larvae collected respectively from BW and FW field habitats, and maintained thereafter in BW and FW respectively, (ii) a self-mating FW colony, termed NFW, established from larvae collected in Nawalapitiya in the central highlands of Sri Lanka (Fig.1) where BW habitats are completely absent, (iii) a self-mating JBW reversal colony (JBWR) established by collecting JBW colony eggs on FW egg-laying surfaces and then hatching and rearing preimaginal stages in FW; and (iv) a self-mating JFW reversal colony (JFWR) established by collecting JFW colony eggs on BW egg-laying surfaces and subsequently hatching and rearing preimaginal stages in BW.

### Cuticle structure in T1 tarsomeres of adult *Aedes aegypti* undergoing preimaginal development in brackish and fresh water

A cuticle covers the external surface of mosquitoes, the gut lumen excluding the midgut and the tracheal lumen [11,15,22,23]. Among other functions, the external cuticle provides surface protection, senses the environment, supports muscle attachment, prevents desiccation in adults, and acts as a barrier for ion and water exchange between preimaginal stages and their aqueous environment [22,23]. Mosquito cuticles are composed of (i) a thin chitin-free epicuticle, not readily visible by transmission electron microscopy (TEM), and composed of highly cross-linked proteins overlaid with a lipid envelope rich in long chain hydrocarbons and waxy esters, and (ii) a wider TEM-visible inner procuticle containing chitin and proteins, and composed of an exocuticle and endocuticle typically synthesized before and after ecdysis respectively [22,23].

We determined cuticle ultrastructure in cross-sections of T1 tarsomeres in the middle-left legs of adult female *Ae. aegypti* three days post-eclosion from the 75^th^ generation (G75) JBW and JFW laboratory colonies by TEM. This was repeated with three days post-eclosion females produced from larvae collected from three separate BW field habitats in the Jaffna peninsula and two days post-eclosion females collected from two different FW field habitats in Nawalapitiya. The tarsi of mosquito legs make the most contact with adulticides when female mosquitoes rest on sprayed surfaces and treated bed nets. TEM images and cuticle thickness measurements are shown in Fig. 2.

**Fig. 2.**
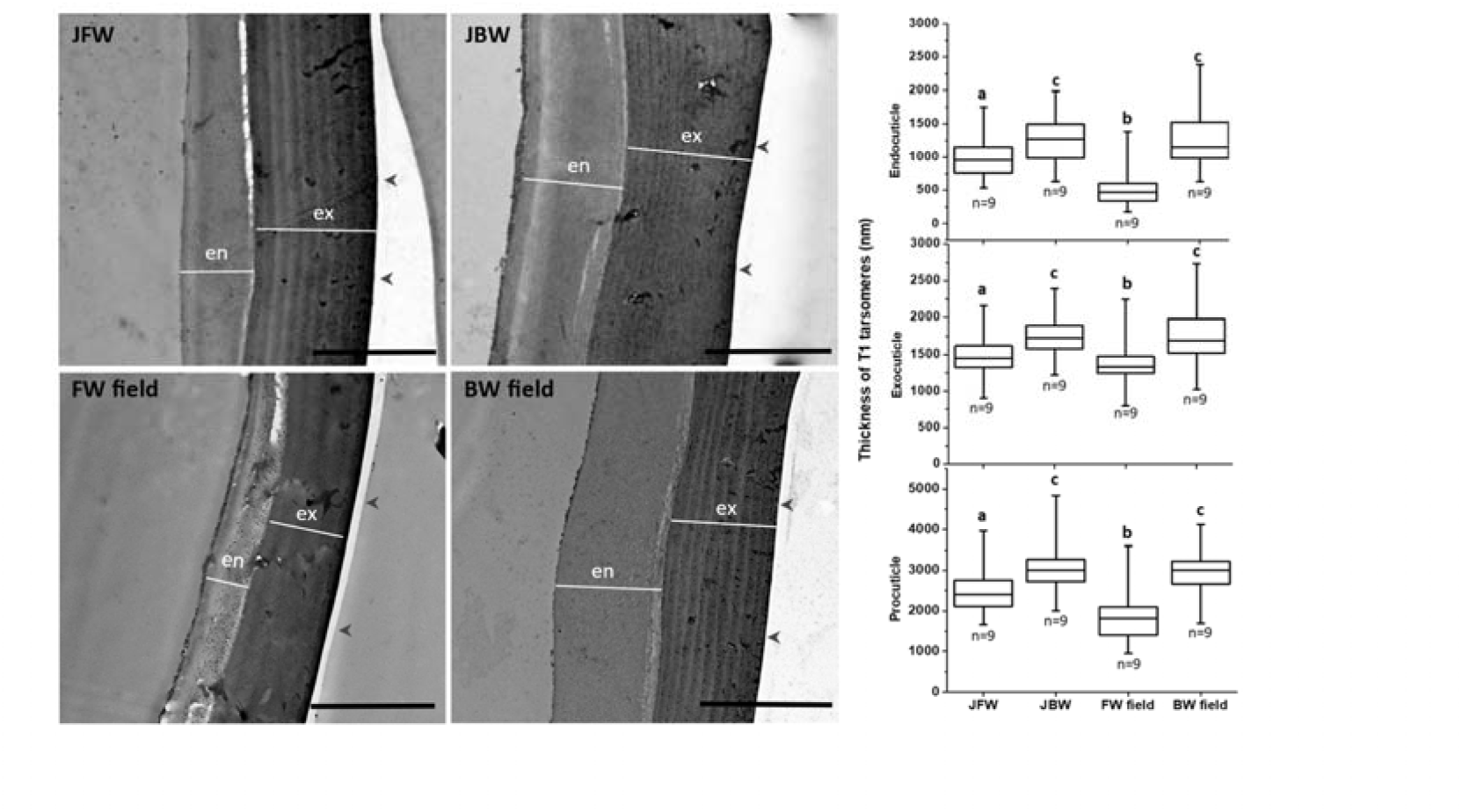
Cuticle ultrastructure and thicknesses of endocuticle, exocuticle and procuticle in T1 tarsomeres of female *Aedes aegypti* emerging from larvae developing in brackish and fresh water. **Legend to Fig. 2** en – endocuticle; ex –exocuticle; n – number of different mosquitoes analysed with 20 measurements in each mosquito; arrowheads indicate the external surface. Box plots show the range (whiskers), median (horizontal line), and 25th and 75th percentiles of measured thicknesses. Values that do not share a letter within each cuticle structure are significantly different (p < 0.05) between the mosquito populations by the Mann-Whitney U test. Statistical test details are shown in Table S1.

Endocuticle, exocuticle and procuticle measurements in the BW and FW field samples were not normally distributed because mosquitoes from different field habitats are typically derived from eggs laid by different females and habitat conditions are never identical. Significance of differences between all mosquito populations was therefore determined by the non-parametric Mann-Whitney U test. The TEM findings showed that the endocuticle, exocuticle and procuticle were significantly thicker in tarsomere T1 of *Ae. aegypti* females produced from larvae developing in BW compared to FW (Fig. 1, Table S1). The endocuticle was particularly thin in FW-field females compared with JFW females, which may partly be due to less complete endocuticle deposition in two rather than three days post-eclosion in tarsi.

### Cuticle ultrastructure in abdomens of female *Aedes aegypti* undergoing preimaginal development in brackish and fresh water

The abdomen and thorax provide large surfaces for absorbing adulticides applied as space sprays indoors and outdoors. TEM was therefore done on mid-6^th^ abdominal segments of a proportion of the same mosquitoes used for T1 tarsomere TEM. TEM images of abdominal cross-sections and the measured cuticle thicknesses are shown in Fig. 3.

**Fig. 3.**
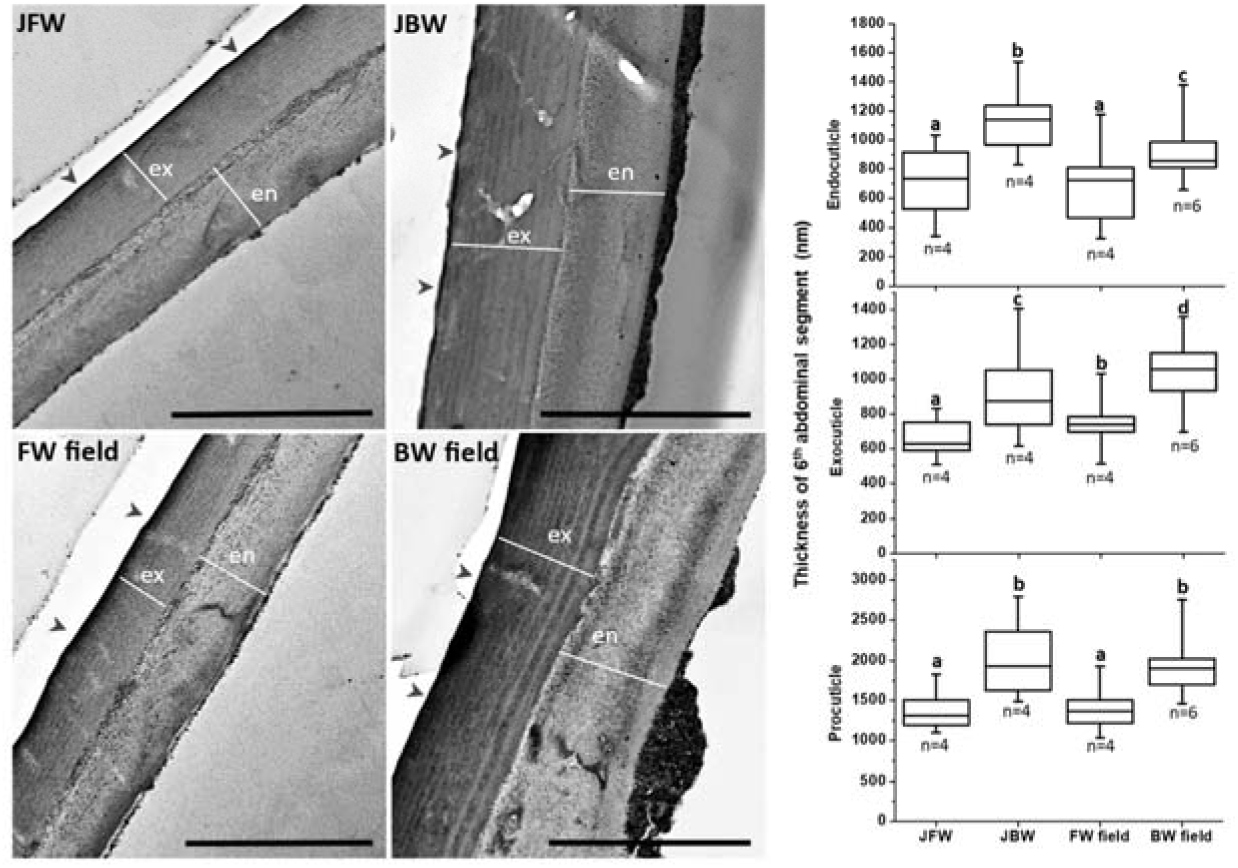
Cuticle ultrastructure and thicknesses of endocuticle, exocuticle and procuticle in the 6^th^ abdominal segments of female *Aedes aegypti* emerging from larvae developing in brackish and fresh water. **Legend to Fig. 3** en – endocuticle; ex –exocuticle; n – number of different mosquitoes analysed with 20 measurements per mosquito; Black scale bars represent 2000 nm. Arrowheads show the external surface. Box plots show the range (whiskers), median (horizontal line), and 25th and 75th percentiles of measured thicknesses. Values that do not share a letter within each cuticle structure are significantly different (p < 0.05) between the mosquito populations by the Mann-Whitney U test. Statistical test details are shown in Table S2.

As in T1 tarsomeres, JBW laboratory and BW field *Ae. aegypti* possessed thicker abdominal endocuticles, exocuticles and procuticles than the corresponding structures in JFW laboratory and FW field *Ae. aegypti.* The endocuticle, exocuticle and procuticle in the 6^th^ abdominal segment were thinner than in T1 tarsomeres from the corresponding mosquito populations, excepting endocuticles in FW field *Ae. aegypti* (Figs. 1 and 2, Table S3), illustrating cuticular differences between the two mosquito structures.

### Insecticide susceptibilities of adult females from *Aedes aegypti* brackish and fresh water laboratory colonies and their reversal colonies

We assessed the susceptibilities to different classes of common adulticides of three-to-five-day-old non-blood fed females from the G63-G72 JBW and JFW, G17-G29 NFW, and G7-G13 JBWR and JFWR colonies. Susceptibilities to permethrin (type 1 pyrethroid lacking a α-cyano group), etofenprox (synthetic type 1 pyrethroid where an ester is replaced with an ether bond), deltamethrin (type 2 pyrethroid containing a α-cyano group), malathion (organophosphate), and propoxur (carbamate) were determined by conventional WHO bioassays for differentiating susceptible and resistant *Ae. aegypti* populations [24,25]. The WHO bioassays use discriminating concentrations of each adulticide determined from a log-probit mortality curve for a reference susceptible mosquito population and then taking twice the concentration of adulticide that produced 100% mortality. Table 1 shows the results of bioassays performed with WHO discriminating concentrations of the adulticides for *Ae. aegypti* [24,25].

**Table 1.**
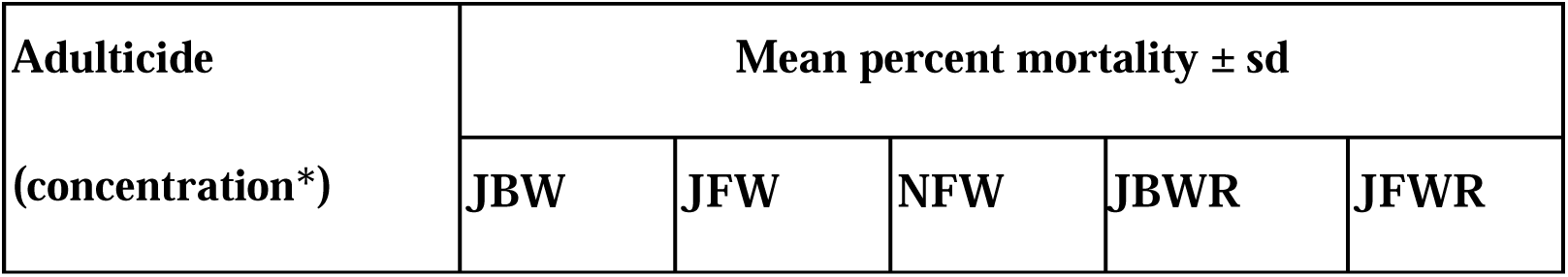
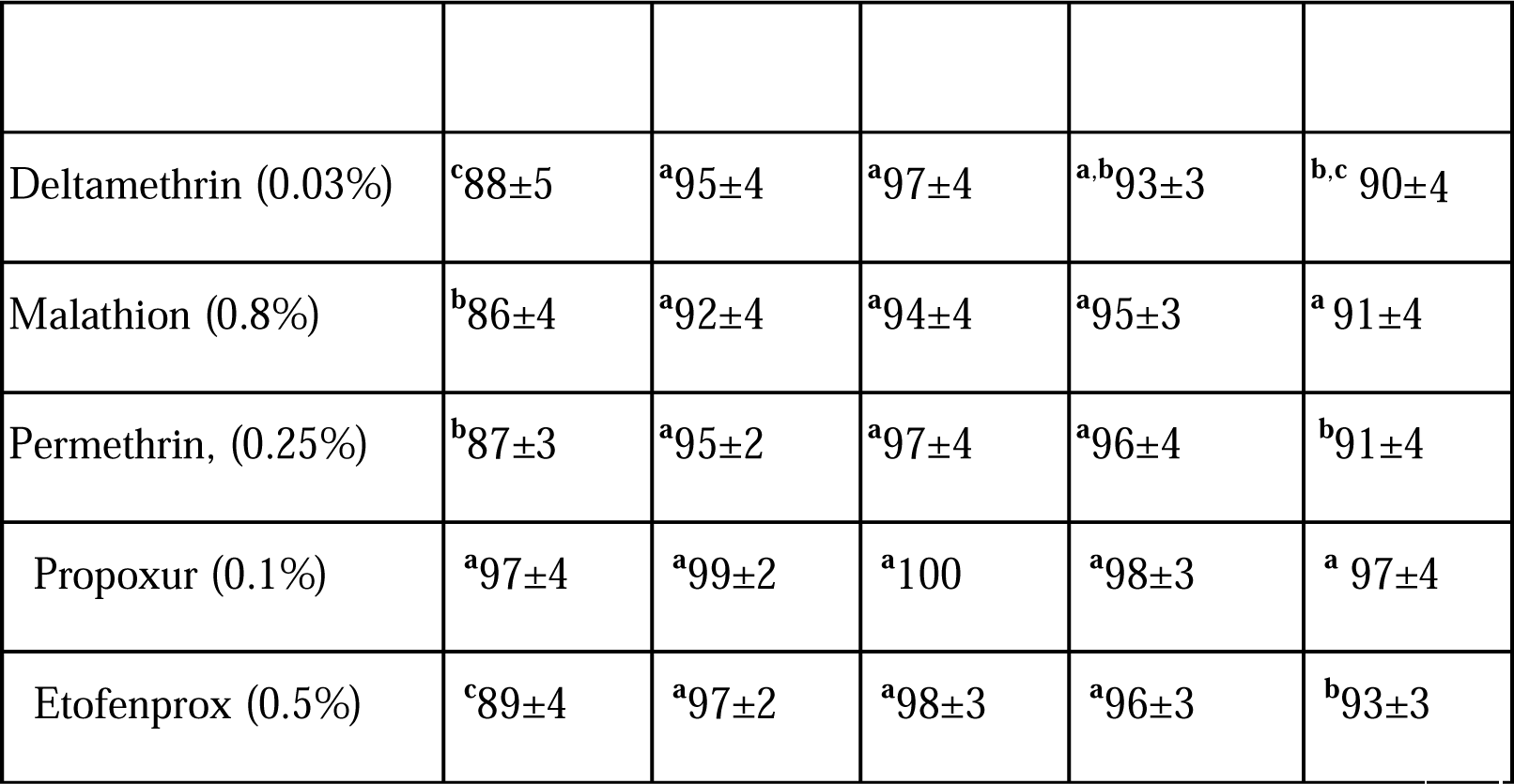
Adulticide susceptibilities of adult female *Aedes aegypti* from laboratory colonies. **Legend to Table 1**. sd – standard deviation. Number of mosquitoes tested per colony per adulticide was 200. Mosquitoes were exposed to adulticide-impregnated filter papers for 1 h and mortalities determined after a 24 h recovery period. Tukey’s method for multiple comparisons in the analysis of variance (ANOVA) was used to compare mean mortalities of test populations. Values that do not share a letter are significantly different (p < 0.05) from others in the same row. *WHO discriminating adulticide concentration [24,25].

The bioassays showed that female JBW *Ae. aegypti* were significantly more resistant to the three pyrethroids and malathion, and tended to be more resistant to propoxur, than either JFW and NFW *Ae. aegypti*. JBW females are categorized as resistant to malathion and all three pyrethroids (<90% mortality) and as possibly resistant to propoxur (90-97% mortality) by WHO criteria [24,25]. In contrast, JFW and NFW were categorized as susceptible to propoxur (≥98% mortality), NFW as susceptible to etofenprox, and both JFW and NFW as only possibly resistant to deltamethrin, malathion and permethrin by the WHO criteria. Resistance of JBW females to the three pyrethroids is particularly significant because pyrethroids are the most common class of adulticides used against *Ae. aegypti*.

The findings from colonies where preimaginal stage salinities were reversed (i.e. from FW to BW for JFWR and from BW to FW for JBWR *Ae. aegypti* respectively) showed that females emerging from reversal colonies tended to alter their adulticide susceptibilities to those corresponding to the new salinities in which the preimaginal stages were reared, i.e. the JFWR approached that of JBW and JBWR was not significantly different from JFW *Ae. aegypti* respectively. The reversal of adulticide susceptibility was evident 7-13 generations after changing preimaginal salinity. The susceptibilities of adult females to the adulticides were therefore determined by the salinity to which their preimaginal stages were exposed.

## Discussion

Widespread use of adulticides selects for increasing adulticide-resistance in *Aedes* field populations, posing a worldwide problem for controlling arboviral diseases [25-29]. Well-established resistance mechanisms include increased activities of adulticide-detoxifying enzymes such as esterases, glutathione-S-transferases and cytochrome P450 monooxygenases (CYPs), as well as mutations in protein targets of adulticides such as acetylcholinesterase and the voltage-gated sodium channel protein (VGSC) [25-31]. There have been very few studies on adulticide resistance in *Ae. aegypti* from the Jaffna peninsula, and these have been restricted to adults developing from FW larval and egg collections [25,32]. The most recent field findings from 2020 with WHO bioassays for permethrin and deltamethrin [32] were compatible with present findings on JFW *Ae. aegypti*, suggesting that the shift to definitive resistance to both pyrethroids in JBW *Ae. aegypti* compared to JFW *Ae. aegypti,* is the result of preimaginal development in BW. Furthermore, the reversal of adulticide susceptibilities in JBWR *Ae. aegypti* showed that preimaginal rearing salinity, and not mutations in genes for acetylcholinesterase, VGSC and detoxifying enzymes, was the determining factor for the greater adulticide resistance in JBW *Ae. aegypti*.

Cuticle changes are another cause of adulticide resistance in mosquitoes and other insects when resistance is selected through exposure to adulticides [30,31,33-41]. A comparison of pyrethroid resistant and susceptible FW laboratory strains of *Ae. aegypti* reported that resistant-strain adults possessed thicker procuticles and higher cuticle polysaccharide content [34]. Similarly, cuticle thickness in T1 tarsomeres was found to correlate with permethrin-resistance in FW laboratory strains of the malaria vector *Anopheles funestus* [35]. Cuticle thickness in femurs [36] and tarsi [37] of FW *Anopheles gambiae* laboratory strains also correlated with resistance to adulticides.

We previously showed that JBW colony larvae were less susceptible to the larvicide temephos than corresponding larvae from the JFW colony [11]. It was established that larval rearing salinity was the determining factor for this difference, and that JBW *Ae. aegypti* larvae also had thicker and less deformable cuticles compared with JFW colony larvae [11,15]. This complements our present finding that the JBW *Ae. aegypti* colony (maintained in 10 g/L salt) and *Ae. aegypti* larvae collected from three BW field habitats (of 6, 6 and 8 g/L salt respectively) give rise to adult females with thicker procuticles, exocuticles and endocuticles in abdomens and T1 tarsomeres of legs than the corresponding structures in females emerging from the JFW colony (maintained in 0 g/L salt) and FW larvae from two field habitats (of 0 g/L salt). The FW field *Ae. aegypti* for the present study were collected from Nawalapitiya where *Ae. aegypti* are not exposed to BW habitats. Hence, the salinity at which preimaginal stages were reared influenced cuticle thickness in adult females as well as larvae in *Ae. aegypti*. Importantly, the thicker abdominal and tarsal procuticles in JBW colony females were associated with greater resistance to common classes of adulticides. A causal relationship between procuticle thickness and greater resistance to adulticides in BW *Ae. aegypti* can be inferred by analogy with findings from adulticide-selected resistance in insects [30,31,33-40], with an additional role for epicuticle changes discussed below. Decreased penetration of adulticides through thicker and remodelled adult cuticles is an underlying mechanism for the resistance induced through adulticide exposure in *An. gambiae* [36] and the oriental fruit fly *Bactrocera dorsalis* [41], and this can also apply to adulticide resistance in female *Ae. aegypti* produced by preimaginal-stage exposure to salinity.

Molecular processes underlying procuticle thickening in larvae and adult procuticles share some features. RNA interference (RNAi) experiments identified a RR-2 family cuticle protein product of the *cpr63* gene [37] and CPLCG5, a member of the CPLCG cuticle protein family [39], as essential for forming the thicker procuticles associated with pyrethroid-selected resistance in adult *Culex pipiens.* Expression of the *cplcg5* gene was controlled by the transcription factor Fushi tarazu-Factor 1 (FTZ-F1), which was upregulated in pyrethroid-resistant adult *Cx. pipiens* [40]. Transcripts for several RR-2 family cuticle proteins as well as FTZ-F1 were also markedly upregulated in JBW L4 exhibiting thicker procuticles and reduced susceptibility to temephos than JFW L4 [11,15].

The lipid-rich envelope of insect epicuticles contains very long chain alkanes formed by elongation of fatty acid acyl CoA, reduction of the elongated fatty acid acyl CoA by a NADH/NADPH-dependent fatty acid acyl CoA reductase to an aldehyde, followed by decarbonylation catalyzed by an insect-specific NADPH-dependent CYP enzyme of the 4G1family characterized first in *Drosophila* [42]. *Drosophila* null-mutants of *cyp4g1* died rapidly after emerging from pupae suggesting that the production of very long-chain alkanes was essential for adults to withstand desiccation [42]. Two homologues of *Drosophila cyp4g1* present in *An. gambiae*, termed *cyp4g16* and *cyp4g17,* encode decarbonylases producing very long chain alkanes, including methyl-branched alkanes [36,37,43]. The two *An. gambiae* decarbonylases were abundant in larval and pupal cells that synthesize epicuticular lipids termed oenocytes [37,43]. *Cyp4g16* and *cyp4g17* are overexpressed in pyrethroid-resistant adult *An. gambiae* laboratory strains, that were also characterized by (i) thicker procuticles and epicuticles, (ii) elevated cuticular hydrocarbon, chitin and chitin-binding cuticular protein content in legs of adult females, and (iii) reduced internalization of topically applied deltamethrin [36,37]. *Aedes aegypti* genes with closest homologies to *An. gambiae cyp4g16* and *cyp4g17* are AAEL004054 (88% coding protein identity to *An. gambiae* CYP4G16) and AAEL006804 (81% protein identity to the available partial sequence of *An. gambiae* CYP4G17) respectively. Their transcripts were more abundant in L4 of JBW than JFW *Ae. aegypti* [16]. Transcripts for chitin synthase, chitin binding proteins, fatty acid synthase, very long chain fatty acid elongase and fatty acyl CoA reductase showed large increases in JBW L4 compared with JFW L4[16]. Increased synthesis of very long-chain cuticular hydrocarbons in the epicuticle, and differential synthesis of chitin and cuticle proteins in the procuticle had been proposed to form thicker and remodelled cuticles in BW-adapted *Ae. aegypti* larvae that minimized the osmotic efflux of water and influx of ions [16]. Epicuticle and procuticle changes thus selected to withstand environmental salinity were proposed to have diminished susceptibility to the larvicide temephos through its reduced uptake [11]. The present findings suggest that aspects of cuticle changes and the underlying molecular mechanisms in BW-adapted *Ae. aegypti* larvae are carried through to adults to reduce susceptibility to common adulticides.

Attrition in FW *Ae. aegypti* transferred to 10 g/L salt BW occurs principally during egg hatching and early larval development [11,14]. Salinity-unrelated environmental stressors causing delayed hatching of eggs have been shown to produce phenotypic changes in adult *Ae. aegypti* [44]. The present findings on adulticide susceptibility in JBWR females and previous findings on temephos susceptibility in JBWR larvae [11] show that these BW-adaptive changes can be reversed within about 13 generations of preimaginal development in FW. The LC_50_ for salt in the L1 to adult transition in the JBWR colony showed incomplete reversal of the LC_50_ after five generations in FW illustrating the slow reversibility of the genetic changes underlying BW adaptation in *Ae. aegypti* [14]. We therefore propose that (i) salinity-induced selection during hatching and early larval development leads to inheritable epigenetic changes expressed in larval and adult *Ae. aegypti* as cuticle changes that respectively reduce susceptibility to larvicides and adulticides, and (ii) such epigenetic changes are slowly reversed on removing the salinity-selection pressure. Similar BW-adaptive changes, including diminished larvicide and adulticide susceptibility, can potentially occur in other FW mosquito vectors recently observed to develop in coastal BW such as *Ae. albopictus*, *An. culicifacies* and *An. stephensi* [6,8,16,17]. BW-developing *Ae. aegypti* and *Ae. albopictus* can be infected with dengue virus and transovarially transmit the virus [45], and therefore can function as virus reservoirs in the absence of vector control measures targeting BW habitats [45]. There is also a potential for FW *Ae. aegypti* adapting to BW to evolve into a new species in isolated coastal locations mirroring, for example, the evolution of the malaria vector *Anopheles merus* from *An. gambiae* in Africa [46].

Our findings highlight the importance of (i) extending mosquito vector control that presently focus on FW preimaginal habitats [1-4] to coastal BW habitats, and (ii) monitoring larvicide and adulticide resistance in BW-developing mosquito vectors and optimizing concentrations of insecticides for use against BW vectors, particularly in the context of global warming raising sea levels [18-21].

## Methods

### *Aedes aegypti* laboratory colonies

Self-mating JBW and JFW laboratory colonies of *Ae. aegypti* initiated with larvae collected from multiple BW and FW habitats in the Jaffna peninsula (Fig. 1) as previously described [6,11,14] were used for experiments. Preimaginal stages of JFW and JBW *Ae. aegypti* were respectively maintained in potable tap water (0 g/L salt) obtained from an artesian well in the centre of the peninsula and sea water diluted with the tap water to yield 10 g/L salt. Because FW *Ae. aegypti* in the Jaffna peninsula were more salinity tolerant than FW mosquitoes from the mainland of Sri Lanka, and there is no barrier to gene flow between JFW and JBW mosquitoes [11,14], we also initiated a laboratory colony termed NFW with *Ae. aegypti* larvae collected from Nawalapitiya in the central hills of mainland Sri Lanka (220 miles from Jaffna Fig. 1), where mosquitoes are not exposed to BW habitats. The NFW colony was maintained in tap water purified by reverse osmosis in the insectary of the Department of Zoology, University of Jaffna. Larvae in colonies were reared in 30 x 25 cm plastic trays containing 1.5 litre water with up to 150 larvae per tray and fed with fish meal powder twice a day. Adults and larvae were maintained at 28–30°C at a relative humidity ∼75% and 12 h dark and light periods. Adult mosquitoes were allowed to feed every three days on Balb/c mice and on 10% glucose pledgets provided at other times. Appropriate 10 g/L salt BW or 0 g/L salt FW surfaces for oviposition were provided in the cages after blood feeding, and eggs harvested two days later as described [11,14]. Stored eggs from early generations of colonies were used to renew the colonies as needed. The 66^th^ generation (G66) of the JBW laboratory colony had a significantly higher 50% lethal concentration (LC_50_) of 16.0 g/L salt than the G66 JFW laboratory colony (LC_50_ of 11.6 g/L) for the L1 to adult transition [11]. Reciprocal crosses between G74 and G75 of JFW and JBW mosquitoes showed them to be reproductively compatible [11]. G17 of the NFW colony had an LC_50_ (11.2 g/L) for the L1 to adult transition which was not significantly different from the JFW colony but significantly lower than the JBW colony [11].

To determine the effects of reversing the larval rearing salinity on adulticide susceptibility, reversal colonies were formed from G63 JBW and JFW colonies as described [11,14]. From the 10 g/L salt-adapted JBW colony, a reversal colony (JBWR) was established by collecting eggs on 0 g/L FW egg-laying surfaces and then hatching and rearing larvae in FW. Similarly, a reversal colony (JFWR) was established from the 0 g/L salt-adapted JFW colony by collecting eggs on 10 g/L salt BW egg-laying surfaces and subsequently hatching and rearing larvae in 10 g/L salt BW.

Mosquito were collected and reared with approval from the Institutional Animal Ethics Committee of the University of Jaffna AERC/2023/03v3.

### *Aedes aegypti* from field habitats

*Aedes* larvae were collected during November and December 2023 from BW habitats in coastal areas of Jaffna city from two different discarded plastic containers with 6 g/L salt water at two sites in Passayoor (9°38’53.628’’N 80°1’49.8’’E; 9°38’5.4168’’N 80°1’56.11’’E) and a discarded plastic yoghurt container with 8 g/L salt water in Columbuthurai (9°39’8.5464’’N 80°2’34.35’’ E) (Fig. 1). *Aedes* larvae were collected in January 2024 at two sites in Nawalapitiya in the central hills of mainland Sri Lanka (Fig. 1) from a discarded plastic flower pot (7°03’17.8"N 80°32’07.3"E) and a discarded plastic bucket (7°03’19.9’’N 80°32’08.8’’E) containing rain water (0 g/L salt). Salinity of larval habitats was measured with a hand-held refracto-salinometer (Atago, Tokyo, Japan). Larvae were collected with 5 ml plastic pipettes and brought to the insectary of the Department of Zoology, University of Jaffna where they were morphologically identified as *Ae. aegypti* using published keys [47] and reared to adulthood in the same water from the habitats from which they were collected. Three-day old BW female adults emerging from BW field habitats and from G75 JFW and JBW laboratory colonies were then processed for transmission electron microscopy (TEM). Logistics associated with larval collection in Nawalapitiya, their maturation to adults in Jaffna, processing and carriage to Switzerland for TEM necessitated using two-day old adult females emerging from larvae collected from FW field habitats in Nawalapitiya for TEM.

### Transmission electron microscopy (TEM)

Tarsomere 1 of the left middle leg and the 6^th^ abdominal segment were dissected from non-blood fed females, washed in 0.1 M Na-cacodylate pH 7.3 buffer (Sigma-Aldrich, MO, USA), and then immersed in cacodylate buffer containing 2% glutaraldehyde (Sigma-Aldrich, MO, USA) for 12-15 h at 4^◦^C. After three washes in cacodylate buffer, post-fixation was performed overnight in 2% osmium tetroxide in cacodylate buffer pH 7.3. Specimens were then transferred to cacodylate buffer pH 7.3 and couriered or hand carried to the University of Bern, Switzerland for further processing and TEM. After three washes in water, specimens were dehydrated sequentially through a graded series of ethanol (30%, 50%, 70%, 90% and 3 × 100%, 2-5 min each). Following dehydration, samples were embedded in Epon812 epoxy resin using plastic embedding molds (Sigma-Aldrich, MO, USA). After polymerization of the resin at 60°C overnight, ultrathin (80 nm) sections were cut using an ultramicrotome (Reichert and Jung, Vienna, Austria). Sections were transferred onto formvar-carbon-coated 200 mesh nickel grids (Plano GmbH, Marburg, Germany), stained with Uranyless® and lead citrate (both from Electron Microscopy Sciences, Hatfield, PA, USA), and imaging performed on a FEI Morgagni transmission electron microscope equipped with a 12 Megapixel Morada digital camera system operating at 80 kV. Tarsomere 1 from nine females each from the JBW and JFW colonies, three females from each of the BW field habitats, four females from one FW field habitat and five from the other, were analyzed by TEM. Abdomens from four females each from the JBW and JFW colonies, two females from each of the BW field habitats, and two females from each FW field habitat were also analyzed by TEM. Thin cross-sections of the mid-region of tarsomere 1 on the left middle leg, while the middle of the 6^th^ abdominal segments was examined in a proportion of the same mosquitoes. Thicknesses of the endocuticle, exocuticle and procuticle from EM images were separately measured referring to the scale bar provided with each image and applying ImageJ [48]. Measurements were taken at 20 different sites distributed across each section of the leg and abdomen. Measured values for the thicknesses of the endocuticle, exocuticle and procuticle were not normally distributed for adults emerging from the larvae collected from either BW field or FW field habitats. Therefore, the significance of differences in cuticle thicknesses between all four mosquito groups was determined by the non-parametric Mann-Whitney U test using Minitab 17 software with a threshold of P<0.05 (Minitab LLC, State College, PA, USA).

### Adulticide susceptibility of JBW, JFW, NFW, JBWR and JFWR of *Aedes aegypti* laboratory colonies

Resistance was determined with adulticide concentrations and procedures recommended by the WHO for differentiating susceptible and resistant *Ae. aegypti* [24] as also previously done in Sri Lanka [25]. Non-blood fed, three-to-five-day old, adult female mosquitoes of G63-G72 JFW and JBW, G17-G29 NFW and G7-G13 JFWR and JBWR, were tested against the recommended WHO discriminating dosage of deltamethrin (0.03%), malathion (0.8%), permethrin (0.25%), propoxur (0.1%) and Etofenprox (0.5%) [24,25]. The assays measure resistance of mosquitoes to tarsal contact with adulticides dispersed in oil and sprayed on filter papers [24,25]. Mosquitoes in batches of 20 were exposed to adulticide impregnated papers for 1 hour at 28 ± 2 [C and 75% relative humidity. The mosquitoes were then removed to different containers and the numbers of dead mosquitoes counted after a 24 h recovery period. Five replicates (20 mosquitoes per replicate) per adulticide were done and experiments performed twice so that a total of 200 mosquitoes were tested with each adulticide. A total of 100 mosquitoes were similarly exposed to filter papers sprayed only with the corresponding solvent oil [24,25] as a control for each adulticide.

Mortality in all control experiments was <5%. Population susceptibility status was as defined by the WHO: susceptible (≥ 98% mortality), possibly resistant (90–97% mortality) and resistant (< 90% mortality) [24]. Tukey’s method for multiple comparisons in the analysis of variance with Minitab 17 software (Minitab LLC, State College, PA, USA) was used to compare the mean percent mortality of the test populations to each adulticide with significance threshold set at p<0.05.

## Acknowledgements

This work was supported in part by the Swiss National Science Foundation through its Program for International Research by Scientific Investigation Teams (IZSTZO: 191762).

## Supplementary information

Table S1. Results of Mann-Whitney U tests for T1 tarsomere cuticle thicknesses

Table S2. Results of Mann-Whitney U tests for abdomen cuticle thicknesses

Table S3. Median values for endocuticle, exocuticle and procuticle thicknesses in T1 tarsomeres and 6th abdominal segment in the different *Aedes aegypti* populations

