## Supplementary Information for "Preimaginal development of *Aedes aegypti* in brackish water produces adult mosquitoes with thicker cuticles and greater insecticide resistance"

**The PDF file includes:**

Tables S1 to S3

**Table S1.** **Results of Mann-Whitney U tests for T1 tarsomere cuticle thicknesses**

| **Samples Compared** | **Exocuticle** | **Endocuticle** | **Procuticle** |
| --- | --- | --- | --- |
| JFW- JBW | *Z*=-9.53945, *p*<0.00001, U=6782 | *Z*= -8.09151, *p*<0.00001, U= 8211 | *Z*=-9.53337, *p*<0.00001, U=6788 |
| JFW- BW field | *Z*=-8.06872, *p*< 0.00001, U= 8234 | *Z*=-7.49998, *p*<0.00001, U=8795 | *Z*=-8.40197, *p*<0.00001, U=7905 |
| JFW-FW field | *Z*=4.46282, *p*< 0.00001, U=11794 | *Z*=13.35401, *p*< 0.00001, U=3016 | *Z*=10.29, *p*<0.00001, U=6041 |
| JBW- BW field | *Z*=0.63103, *p=*0.5287, U=15577 | *Z*=0.90705, *p=*0.36282, U=15304 | *Z*=0.54848, *p=*0.58232, U=15658 |
| JBW-FW field | *Z*=11.70248, *p*< 0.00001, U= 4646 | *Z* =15.21217, *p*<0.00001, U= 1181 | *Z*=13.97289, *p*<0.00001, U=2405 |
| BW field-FW field | *Z*=10.55285, *p*<0 .00001, U= 5781 | *Z*=15.07441, *p*< 0.00001, U= 1317 | *Z*=13.33932, *p*<0.00001, U=3030 |

**Table S2.** **Results of Mann-Whitney U tests for abdomen cuticle thicknesses**

| **Samples**  **compared** | **Exocuticle** | **Endocuticle** | **Procuticle** |
| --- | --- | --- | --- |
| JFW-JBW | *Z*=-8.90864, *p*<0.00001, U= 589 | *Z*=-9.33863, *p*<0.00001, U=463 | *Z*=-9.3574, *p*<0.00001, U=458 |
| JFW-BW field | *Z*=11.45764, *p*<0.00001, U=205 | *Z*=5.70201, *p*<0.00001, U=2513 | *Z*=10.57485, *p*<0.00001, U=559 |
| JFW-FW field | *Z*=5.45848, *p*< 0.00001, U=1600 | *Z*=-0.63816, *p*=0.52218, U=3013 | *Z*=0.60915, *p*=0.54186, U=3021 |
| BW-BW field | Z=5.17707, *p*<0.00001, U=2724 | *Z*=-8.0499, *p*<0.00001, U=1572 | *Z*=-1.0025, *p*=0.31732, U=4398 |
| BW-FW field | *Z*=5.74856, *p<*0.00001, U=1515 | *Z*=9.43248, *p*<0.00001, U=436 | *Z*=9.54168, *p*<0.00001, U=404 |
| BW field-FW field | *Z*=10.60976, *p*< 0.00001, U=545 | *Z*=6.56611, *p*< 0.00001, U=2167 | Z=10.61849, *p*< 0.00001, U=542 |

**Table S3. Median values for endocuticle, exocuticle and procuticle thicknesses in T1 tarsomeres and 6^th^ abdominal segment in the different *Aedes aegypti* populations**

**(a) Cuticle thickness in T1 tarsomeres**

| Mosquito population | Exocuticle thickness  median in nm | Endocuticle thickness  median in nm | Procuticle thickness  median in nm |
| --- | --- | --- | --- |
| JFW (n=9) | 1452 | 958 | 2397 |
| FW field (n=9) | 1331 | 475 | 1824 |
| JBW (n=9) | 1722 | 1272 | 3010 |
| BW field (n=9) | 1691 | 1146 | 3014 |

**(b) Cuticle thickness in mid-sixth abdominal segment**

| Mosquito population | Exocuticle thickness  median in nm | Endocuticle thickness  median in nm | Procuticle thickness  median in nm |
| --- | --- | --- | --- |
| JFW (n=4) | 628 | 732 | 1306 |
| FW field (n=4) | 740 | 721 | 1369 |
| JBW (n=4) | 876 | 1142 | 1925 |
| BW field (n=6) | 1059 | 857 | 1902 |

**Legend to Table S3**. n – number of different mosquitoes analysed with 20 measurements per mosquito
